# Spatio-temporal evolution of urban thermal environment and its driving factors: Case study of Nanjing, China

**DOI:** 10.1101/2021.01.13.426518

**Authors:** Zhang Menghan, Dong Suocheng, Cheng Hao, Li Fujia

## Abstract

In recent years, with the rapid urbanization, the urban underlying surface has changed dramatically. Various urban eco-environmental problems have emerged in the world, among which the urban heat island effect has become one of the most obvious urban eco-environmental problems. In this study, Nanjing, China is chosen as the study area. Based on the Landsat8 remote sensing image data in Nanjing from 2014 to 2018, the land surface temperature is retrieved, the spatiotemporal variation track and characteristics of the thermal environment pattern are systematically depicted, and its driving factors are revealed. The results show that: in the past five years, the spatial pattern of heat field in Nanjing has changed from scattered distribution in the periphery of the city to centralized in the center of the city, and the heat island intensity has increased year by year. The changes of administrative divisions, the layout of transportation trunk lines, the transfer of industrial centers, and ecological construction projects are important driving factors for the evolution of land surface thermal environment pattern of these regions. The research results will provide scientific and technological support for similar cities with typical heat island effect in the world to make urban planning and development decisions, and to govern and improve urban ecological environment.

## Introduction

In recent years, with the continuous acceleration of urbanization, urban areas have become a hot spot for cross-scale research on the driving factors of environmental change [1]. In the process of rapid urbanization, many ecological and environmental problems appear in the developing countries represented by China, which directly lead to the drastic changes of urban surface thermal environment. Whether the urban surface thermal environment is good or not is one of the important indexes to measure the urban ecological environment [2]. Urban areas generally exhibit higher air and surface temperatures than surrounding rural areas [3], and tends to raise local temperatures, resulting in an *urban heat island (UHI)* [4]. UHI is a kind of concentrated reflection and embodiment of urban space surface thermal environment to some extent [2], and the phenomenon of UHI has long attracted great interest from scientists and urban planners.

Academia’s attention to the UHI has been more than 200 years of history [5–6]. In recent years, with the rapid development of urbanization and ecological protection awareness of the world, UHI has been more and more common concern of scholars at home and abroad. Scholars from all over the world have carried out a lot of research on the characteristics [7–9], influencing factors [10–12], simulation and countermeasures [13]and other aspects of UHI. Shastri, H et al for the first time analyzed the day and seasonal characteristics of *surface urban heat island intensity (SUHII)* in urban centers of India and found that SUHII in most urban areas was negative during the pre-monsoon summer. Wu, X et al taking Dalian, a coastal city in China, as an example, revealed the dynamic mechanism of the UHI effect for different seasons using the cubist regression tree algorithm and suggested that the UHI effect only existed in spring and summer with moderate value, and no obvious UHIs can be found in autumn and winter. Ngarambe, J et al explored the synergies between UHI and HW in Seoul city, and showed that UHI is more intense during HW periods than non-heat wave (NHW) periods and the synergies were relatively more intense in densely built areas and under low wind speed conditions. Yang, Q et al applied temperature and land-cover classification datasets based on satellite observations to analyze that that SUHII and its correlations with ΔLMs were profoundly influenced by seasonal, diurnal, and climatic factors in 332 cities/city agglomerations distributed in different climatic zones of China. Manoli, G et al introduced a coarse-grained model that links population, background climate, and analyzed summertime differences between urban and rural surface temperatures (ΔTs) worldwide, showing that urban–rural differences in evapotranspiration and convection efficiency are the main determinants of warming. Sangiorgio, V et al established a multi-parameter calibration index for comprehensive evaluation of UHI. It was found that urban albedo and the existence of green plants were the most important factors affecting the potential absolute maximum *urban heat island intensity (UHII)* in urban districts. Li, Y et al used the surface and vegetation characteristics of the region around Berlin (Germany), and run the urban climate model driven by the same lateral climate conditions in order to simulate the urban climate of various generated cities under the same weather conditions.

In the context of rapid urbanization, especially in developing regions [14], people are also increasingly concerned about the risks posed by the urban heat island effect on urban residents [11,15–17]. Mora, C et al conducted a global analysis of documented lethal heat events, and found 783 cases of excess human mortality from 164 cities in 36 countries associated with heat. Werbin ZR et al pointed out that the increasing exposure of urban heat island effect led to the increasing high mortality and morbidity associated with high temperature, which posed an urgent threat to urban public health. Manoli, G et al also pointed out that UHIs exacerbate the risk of heat-related mortality associated with global climate change. Urban surface thermal environment is not only directly related to the quality of urban living environment and residents’ health, but also has a far-reaching impact on urban energy and water consumption, ecosystem process evolution, biological phenology and sustainable economic development [1,18–21]. Therefore, it is imperative to study the urban heat island effect and its influencing factors, and at the same time, it becomes a particularly urgent problem to put forward countermeasures to improve a series of risks and negative effects caused by it.

In this paper, Nanjing, China, with typical heat island effect, is selected as the case study. By using Landsat8 remote sensing image data from 2014 to 2018, the surface temperature and surface feature parameters of the third phase are obtained, on the basis of which, the thermal environment pattern of Nanjing city and the characteristics of its spatio-temporal evolution are illustrated and the driving factors and change of the urban thermal environment pattern under different space conditions are further explored. The research results will provide scientific and technological support for similar cities with typical heat island effect around the world to make urban planning and development decisions, and to govern and improve urban ecological environment.

## Materials and methods

### Case study

Nanjing is located in the southwest corner of Jiangsu Province (Fig 1). It is located in the lower reaches of the Yangtze River and adjacent to the Yangtze River and the East China Sea. Its geographical coordinates are 31 ° 14 ″ ~ 32 ° 37 ″ N and 118 ° 22 ″ ~ 119 ° 14 ″ W. Nanjing is located in the hilly area of Nanjing-Zhenjiang-Yangzhou, with low mountains and gentle hills. The forest coverage rate of the whole city is 27.1%, and the water area is more than 11%. There are Qinhuai River, Jinchuan River, Xuanwu Lake, Mochou Lake and other rivers and lakes. The Yangtze river passes through the city, and the total length of its coastline is nearly 200 km.

**Fig 1.**
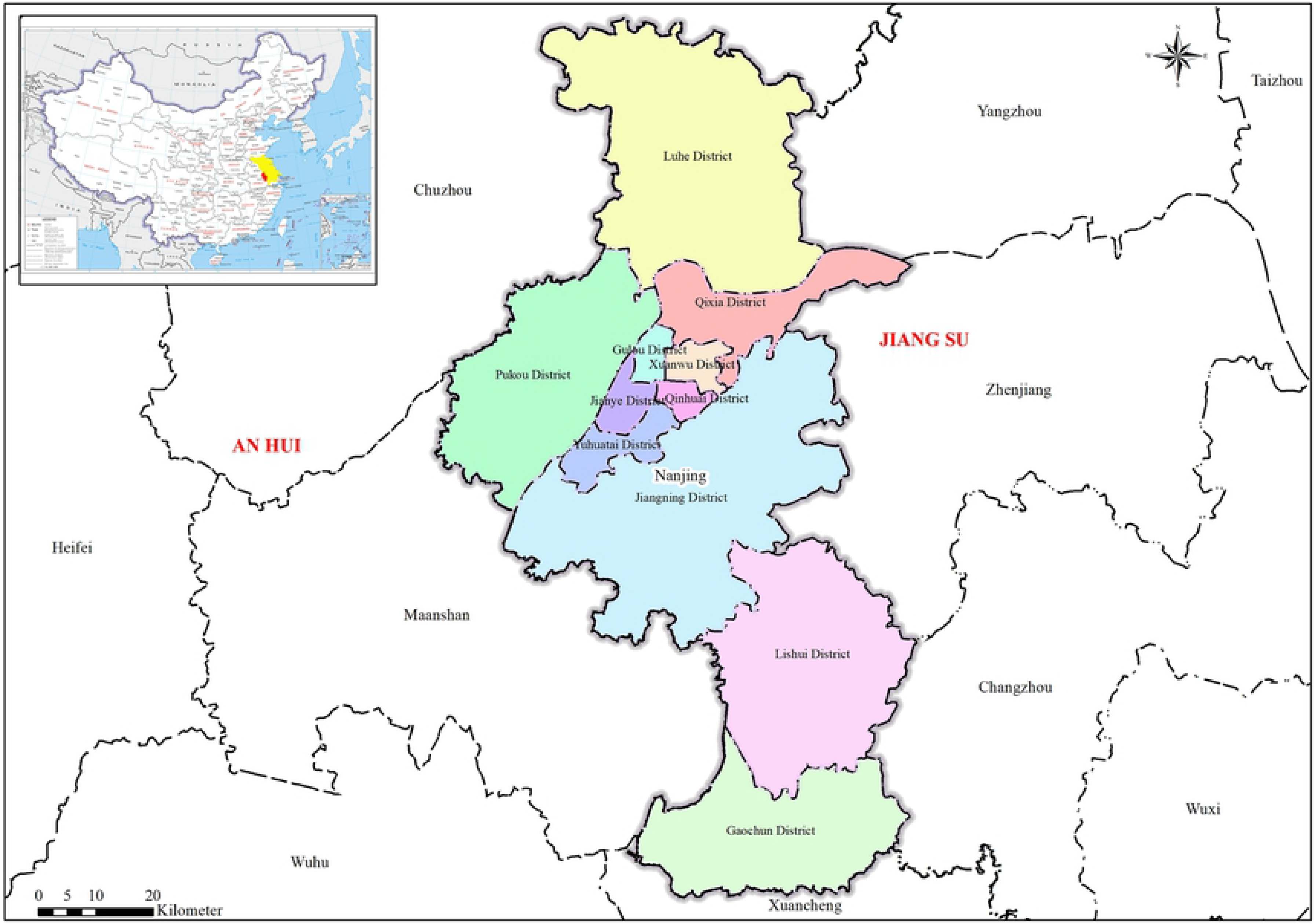
Location Map of the Study Area.

Nanjing has an urban construction history of nearly 2600 years and a capital city history of nearly 500 years. It can be traced back to the city of Yecheng, which was put here by the Wu state in the Spring and Autumn and the Warring States Period. Nanjing has a large population. By 2018, there were 8,436,200 permanent residents in Nanjing, with an urbanization rate of 82.5 percent. Nanjing is the vice-provincial city and also the capital of Jiangsu Province and enjoys a high political status. What’s more, according to the Notice on the Adjustment of Standards for the Division of City Size issued by the State Council (Document No. 51 of 2014), Nanjing is classified as a megacity with a permanent urban population of more than 5 million. In addition, Nanjing is also an important gateway city planned and positioned by the State Council for the Yangtze River Delta to radiate and drive the development of the central and western regions. It is also an important node city at the strategic intersection of the eastern coastal economic belt and the Yangtze River Economic Belt, and has an important strategic position.

### Data

The collected remote sensing image data were less cloud and cloudless, with good quality, and fully cover the three phases of Landsat8OLI with strip number as 120 and row number as 38 in Nanjing. The imaging time was November 17, 2014, March 28, 2016 and April 19, 2018, respectively (Table 1). This research used the 4th, 5th and 10th bands of OLI for land surface temperature retrieval. Other auxiliary information included the remote sensing land use remote sensing monitoring data and administrative division data of Nanjing in 2018 (Table 2).

**Table 1.** The detailed information of remote sensing data.

| Data acquisition time | Remote sensor | Data number | Longitude | Latitude |
| --- | --- | --- | --- | --- |
| 2014/11/17 | Landsat8OLI_TIRS | LC81200382014322LGN00 | 118.846 | 31.738 |
| 2016/3/28 | Landsat8OLI_TIRS | LC81200382016088LGN00 | 118.838 | 31.738 |
| 2018/4/19 | Landsat8OLI_TIRS | LC08_L1TP_120038_20180419_20180501_01_T1 | 118.831 | 31.738 |

**Table 2.** Research data and its sources.

| Research data | Data sources |
| --- | --- |
| Landsat8OLI | Geospatial Data Cloud (http://www.gscloud.cn/) |
| Land use monitoring data of Nanjing in 2018 | Resource and Environment Science and Data Center (http://www.resdc.cn/lds.aspx?tdsourcetag=s_pctim_aiomsg) |
| Administrative division data of Nanjing | Geospatial Data Cloud (http://www.gscloud.cn/) |

**Table 4.** Results of atmospheric parameters of remote sensing images in each year.

| Data acquisition time | Atmospheric upward radiance ($W/m^2/sr/\mu m$) | Atmospheric downward radiance ($W/m^2/sr/\mu m$) | Atmospheric transmittance |
| --- | --- | --- | --- |
| 2014/11/17 | 0.33 | 0.58 | 0.95 |
| 2016/3/28 | 0.45 | 0.80 | 0.93 |
| 2018/4/19 | 0.84 | 1.45 | 0.90 |

**Table 5.** Surface temperature statistics (°C) in each year.

| Year | Minimum | Maximum | Mean | Standard deviation |
| --- | --- | --- | --- | --- |
| 2014 | 1.4 | 25.6 | 14.7 | 1.4 |
| 2016 | 7.8 | 30.8 | 22.4 | 2.7 |
| 2018 | 18.4 | 42.0 | 29.6 | 3.3 |

In order to further explore the reasons of land surface thermal environment changes in Nanjing, this research conducted the on-site investigation and survey on the key regions with obvious changes in the heat island intensity, and mainly investigated the current situations of Gaochun District and Lishui District, construction and surrounding development conditions of Jiangbei Avenue Expressway, construction of Nanjing Jiangbei New Materials High-Tech Park, and conditions of ecological restoration and protective development of Laoshan National Forest Park and Zijinshan National Forest Park in Nanjing (S1 Table).

### Research methods

#### Atmospheric correction method

This study was based on the atmospheric correction method (also known as the radiation transfer equation method, RTE), and used the three phases of Landsat8OLI_TIRS remote sensing images for land surface temperature retrieval. Its basic principle was firstly estimating the influence of the atmosphere on the surface thermal radiation, then subtracting such atmospheric influence from the total thermal radiation observed by satellite sensors, thus obtaining the surface thermal radiation intensity, and transforming the thermal radiation intensity into corresponding surface temperature.

The radiation transfer equation of the satellite sensor receiving the thermal infrared radiance value *L*_*λ*_ was:

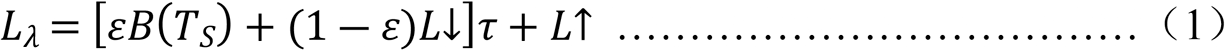

In formula (1), ε was the land surface emissivity, *T*_*S*_ was the real surface temperature (K), *B*(*T*_*S*_) was the blackbody thermal radiance, *τ* was the atmospheric transmission in thermal infrared band. Thus the radiance *B*(*T*_*S*_) of the blackbody with temperature *T* in the thermal infrared band could be calculated by formula (2).

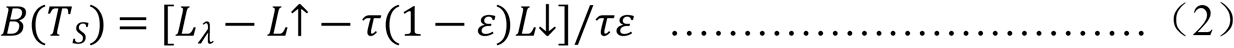

In formula (2), *T*_*S*_ can be obtained by the functions of Planck formula.

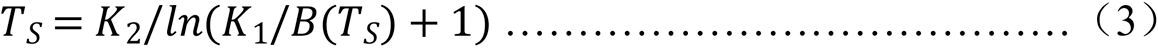

In formula (3), for TIRS Band10, K1=774.89W/(m2*μm*sr), K2=1321.08K.

Atmospheric profile parameters could be obtained from the website which was provided by NASA (http://atmcorr.gsfc.nasa.gov/).The values of atmospheric upward radiance, downward radiance and transmittance were as follows:

#### Urban heat island intensity

The southern regions in China had the common weather of rainy days and cloudy days, the images of the same period in many years were hardly obtained, so theoretically the remote sensing images of the same time phase were also hardly obtained. The research of heat island effect mainly focused on analyzing the spatial distribution characteristics of the relative strength of urban underlying surface temperature, and the difference of seasons only changed the size of land surface temperature instead of its spatial distribution.

Therefore, in order to reflect and display the urban land surface thermal environment in a more concentrated way, the concept of *urban heat island intensity (UHII)* was introduced. The UHII referred to the difference between the average temperature in the center of a city and that in the surrounding suburbs (villages), which was used to indicate the intensity of urban heat island effect. The calculation formula of UHII was as follows:

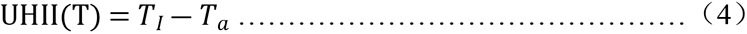

In formula (4), UHII(T) was the urban heat island intensity, *T*_*I*_ was the surface temperature at a certain point in urban area, *T*_*a*_ was the average temperature in the suburbs, and land surface temperature of farmland was used instead of retrieved surface temperature.

## Results

### Spatial-temporal distribution pattern of Nanjing land surface thermal environment

The 3 spatial distribution diagrams of land surface thermal environment (Fig 2) showed that, from 2014 to 2018, the spatial distribution of Nanjing urban land surface thermal environment had the below features:

1. In 2014, the maximum surface temperature of Nanjing was 25.6°C, the minimum temperature was 1.4°C, and the average temperature was 14.7°C. The northern regions of Nanjing had higher temperature, especially the temperature of riverside belts on the north of Tangshan was particularly high, and the administrative centers of various districts and minor industrial zones formed many scattered land surface temperature high-value regions.
2. In 2016, the maximum surface temperature of Nanjing was 30.8°C, the minimum temperature was 7.8°C, and the average temperature was 22.4°C. The west-central regions of Nanjing had higher temperature, and the northwest corner had much higher temperature than the surrounding regions. In view of the overall spatial distribution features, the urban land surface thermal environment started showing its distribution along main roads, and formed many strip land surface temperature high-value regions. The temperature of rivers, lakes and mountain regions was relatively lower.
3. In 2018, the maximum surface temperature of Nanjing was 42.0°C, the minimum temperature was 18.4°C, and the average temperature was 29.6°C. The Nanjing urban land surface thermal environment started showing the distribution features of concentrated block masses, and the high-value regions were more concentrated, which were mainly distributed around the main urban areas in the southern regions of Yangtze River and Jiangbei New Area.

**Fig 2.**
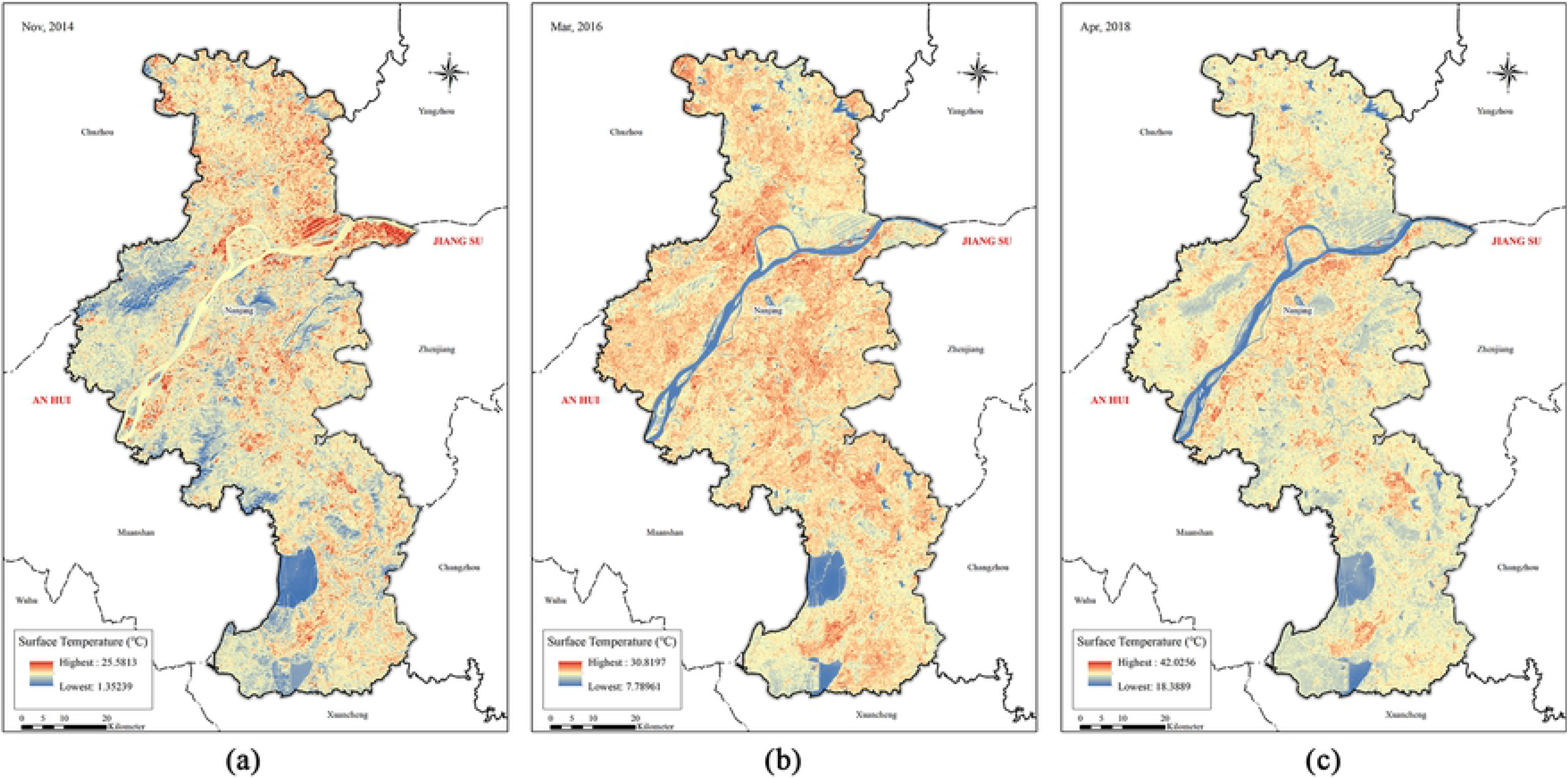
A spatial distribution map of heat field in Nanjing in 2014(a),2016(b),2018(c).

### Comparison and analysis of multi-phase Nanjing land surface thermal environment distribution pattern

The Nanjing urban land surface thermal environment spatial pattern was reflected and displayed in a concentrated way relying on the heat island intensity. The analysis found that, from 2014 to 2018, the Nanjing UHII spatial pattern had the below features (Fig 3):

1. In 2014, Nanjing heat islands appeared in the form of scattered distribution, and were mostly concentrated around the riverside belts on the north of Tangshan. The heat island effect of the whole Nanjing city was mainly concentrated around the northern regions and northern riverside belts, but was relatively less in the southern regions.
2. In 2016, the UHIs still showed the scattered distribution, but had a wider distribution area from the riverside regions on the north of Tangshan to the whole Nanjing city, and showed many heat islands in strip distribution along main roads. The overall spatial distribution showed the trend of gradually moving to the southwest direction and being accompanied with the obvious diffusion and dilution status.
3. In 2018, the UHIs covered the main urban areas and Jiangbei New Area of Nanjing, and started showing the distribution features of concentrated block masses, and wholly showed the distribution features of being high and concentrated in central regions, being low and scattered in surrounding regions. Meanwhile, the heat island intensity was increased obviously, the heat island effect was significant, and the land surface temperature in central urban areas was obviously higher than that of outer suburbs. Besides the main urban areas and Jiangbei New Area, the heat islands of central areas of various towns in southern regions were also strengthened.

**Fig 3.**
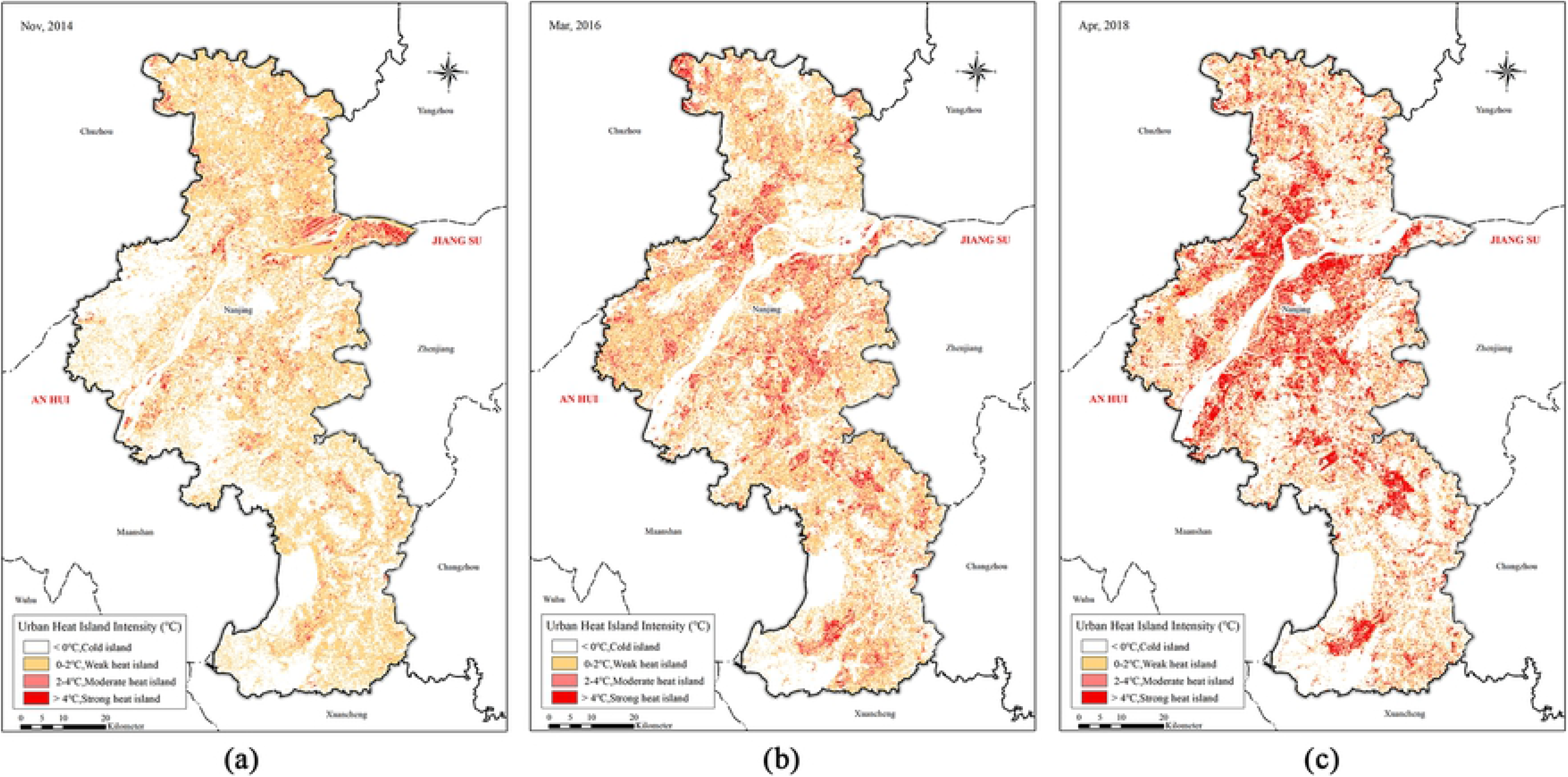
A spatial distribution map of heat island intensity in Nanjing in 2014(a),2016(b),2018(c).

From 2014 to 2018, the spatial pattern of Nanjing UHII (Fig 4) had obvious changes. The change volume of heat island intensity had the average value of 0.01°C, maximum value of 25.6°C, and minimum value of −13.7°C, which were respectively located in Yaohuamen regions of Qixia District and on river surface of northern Yangtze River. The changes of Nanjing heat island intensity wholly showed the trend of “ascending in urban areas and riverside regions, and descending in suburbs”. Thereinto, the suburbs in southern Nanjing had two “heat islands”, and the urban areas had two “cold islands”. The features were mainly summarized as below:

1. The heat island intensity of urban areas and riverside regions ascended obviously. The regions of heat island intensity ascending obviously were concentrated around the main urban areas in the southern regions of Yangtze River, Jiangbei New Area and the upstream and downstream southern bank regions of Yangtze River Nanjing Section. Thereinto, in the northern regions of Yangtze River, the regions of heat island intensity ascending obviously showed the distinct strip distribution, and were divided into two-way extension from northeast to southwest direction.
2. The heat island intensity of suburbs and water regions descended obviously. The regions of heat island intensity descending obviously were concentrated around the Donggou Town of Luhe District in suburbs and riverside belts on the north of Tangshan. Meanwhile, the whole Yangtze River Nanjing Section water regions, the southern Shijiu Lake and Gucheng Lake water regions all belonged to the regions of heat island intensity descending obviously.
3. The heat island intensity of outer suburbs kept a stable status. Except for Shijiu Lake and Gucheng Lake water regions in the southern suburbs of Nanjing, the heat island intensity of outer suburbs in the both south and north of Yangtze River did not have obvious changes, and the difference between the land surface temperature of outer suburbs and that of urban areas always kept a stable status.

**Fig 4.**
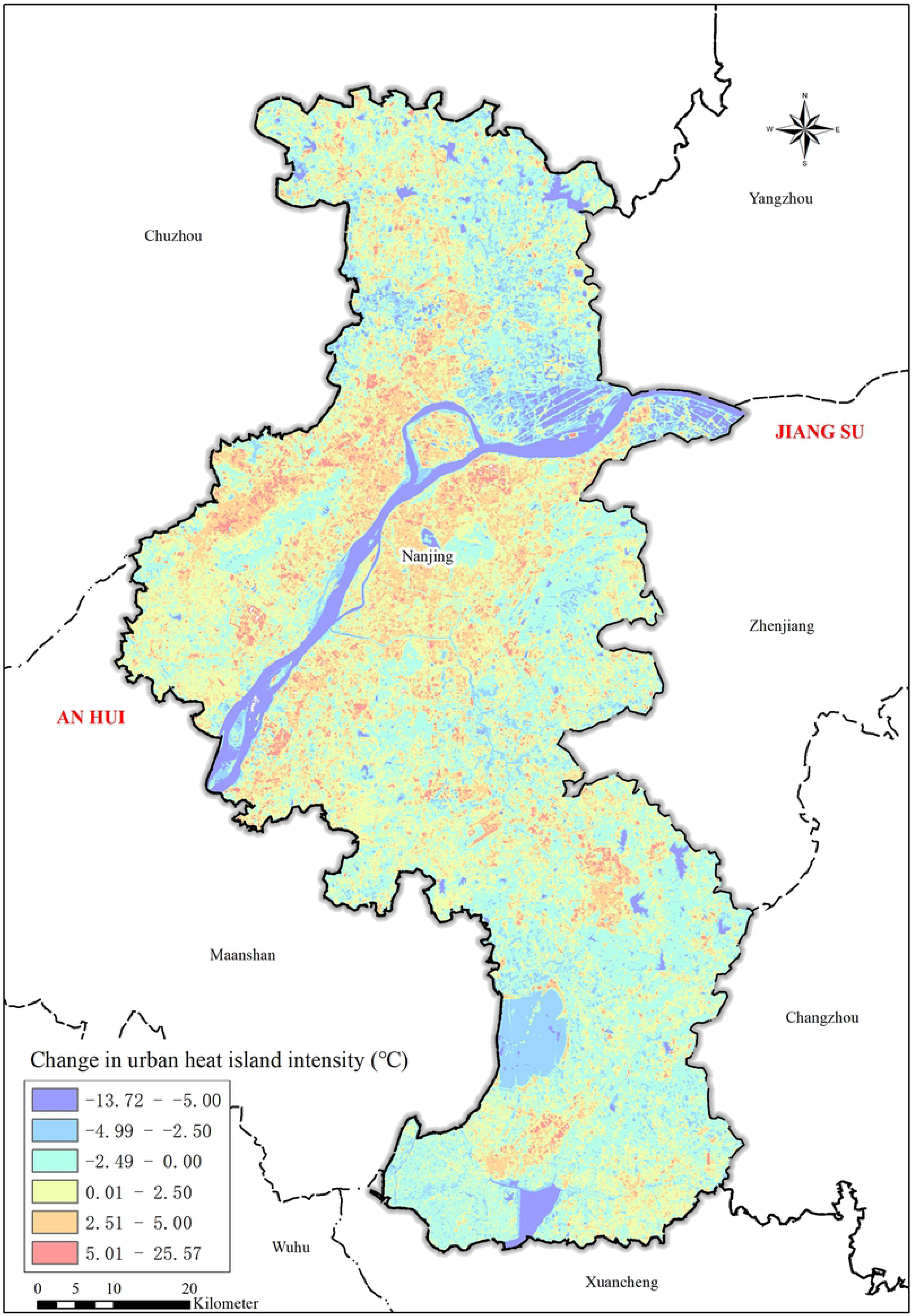
A spatial and temporal variation map of heat island intensity in Nanjing city from 2014 to 2018.

### Driving factors of Nanjing land surface thermal environment formation

In order to further explore the driving factors of changes of Nanjing heat island intensity, this research conducted the on-site investigation and survey on the above regions with obvious changes in heat island intensity, and mainly investigated the changes of such regions in the research periods. According to the investigation and survey results, this research obtained the below conclusions:

1. The changes of administrative division caused transfer of heat island centers This research conducted the investigation and survey aiming at two obvious “heat islands” which appeared at outer suburbs in the southern Nanjing, and found two “heat islands” which were respectively the administrative centers of Gaochun District and Lishui District. According to the file of *Reply of the State Council on Agreeing with Jiangsu Province Adjusting Some Administrative Division of Nanjing (GH [2013] No. 24)*, Nanjing was approved to adjust some administrative division in 2013, cancel the former Lishui County and Gaochun County and establish the Lishui District and Gaochun District, and merge them into the jurisdiction of Nanjing. After the cancellation of counties and establishment of districts, following the continuous development of new town construction, industrial transformation and urban economy of such two districts and continuous deep-going of urban-rural integration process, the Gaochun District and Lishui District which were originally far away from the main urban areas of Nanjing and had backward development caught the development opportunities, followed the trend, and gradually entered the accelerating development stage of urbanization. Therefore, the obvious urban heat island effect appeared in surrounding regions of their administrative centers, Yongyang Sub-district of Lishui District and Chunxi Sub-district of Gaochun District, and had influences on the spatial distribution of Nanjing land surface thermal environment.
2. The surrounding regions of core transportation lines were developed quickly, and prompted the linear gathering of land surface thermal environment. Aiming at the regions of heat island intensity ascending obviously in northern regions of Yangtze River, this research analyzed and found such regions showed the obvious distribution along transportation roads, and mainly extended along two parallel northeast-southwest direction expressways (Jiangbei Section of Nanjing Around-city Expressway and Jiangbei Avenue Expressway). Nanjing undertook the Summer Youth Olympic Games 2014. As the transportation guarantee project serving the Summer Youth Olympic Games 2014, Jiangbei Avenue Expressway completed the main line construction work in 2014, and continued making multiple stages of transformation works. Under this influence, the road transportation network of northern regions of Yangtze River became gradually mature, realized series connection of the whole northern regions of Yangtze River sector, formed joint forces with subways, completed the road network structure of the whole Jiangbei New Area, shorten the distance from the northern regions of Yangtze River to the main urban area in the southern regions of Yangtze River, and attracted more enterprises to settle down in Jiangbei New Area. The increasingly frequent construction and commuting behaviors significantly enhanced the heat island intensity of such regions. In the regions of heat island intensity ascending obviously, an area where two expressways approach each other appears to be connected into patches. The main reason accounting for this phenomenon was that the Nanjing Jiangbei New Materials High-Tech Park here which was established in October 2001 played the role of migration place of chemical enterprises in the southern regions of Yangtze River during the process of the projects of closure and relocation of chemical pollution enterprises which were vigorously implemented by the Nanjing Municipal government. Following the continuous increase of the quantity and scale of migration chemical enterprises, the heat island effect around the Nanjing Jiangbei New Materials High-Tech Park ascended obviously. The regions between the Jiangbei Avenue Expressway and the Yangtze River bank forming the regions of heat island intensity ascending obviously with concentrated connection in patches was mainly because that, following the development of Yangtze River shipping, a large quantity of transportation facilities was distributed along the Nanjing riverside belts, and their quantity and scale were increased year by year. Aiming at the regions of heat island intensity ascending obviously in the southern regions of Yangtze River, this research analyzed and found that they were concentrated around the urban areas with the Nanjing Around-city Expressway as boundaries. The regions out of the Nanjing Around-city Expressway circle lines were mainly extended to periphery along Changchun-Shenzhen Expressway and Nanjing-Xuancheng Expressway, and had the conditions of connection in patches close to the administrative center of Lishui District. Meanwhile, similar with the northern regions of Yangtze River, the regions between the roads and the Yangtze River bank also formed the regions of heat island intensity ascending obviously with concentrated connection in patches, covered the whole southern bank of Yangtze River Nanjing Section, and had a larger scale of concentrated connection in patches than that of the northern regions of Yangtze River.
3. The transfer of urban industrial centers prompted “heat islands” being transformed into “cold islands”. Aiming at the regions of heat island intensity descending obviously in the southern regions of Yangtze River, this research analyzed and found that these regions were caused by the projects of closure and relocation of chemical pollution enterprises which were vigorously implemented by the Nanjing Municipal government. According to the requirements of “Blue Sky Action Plan (2010-2015) which was started by the Nanjing Municipal Government in 2010, the 66 medium and small chemical enterprises within the range on west of around-city roads in Yanqi regions should be closed, and 3 chemical enterprises should be relocated to Nanjing Jiangbei New Materials High-Tech Park; the chemical enterprises around the SINOPEC Jinling Company should be renovated, more than 20 small chemical enterprises should be closed, transformed and governed. The goal of this plan is to basically realize that there is no chemical enterprise within the around-city roads in the southern regions of Yangtze River. Under this influence, the heat island effect of such regions showed descending to certain extent.
4. The urban ecological project construction prompted the forming of “cold island” regions. Aiming at the two “cold islands” in the urban areas, this research conducted the investigation and survey, and found such “two islands” were respectively the Laoshan National Forest Park of Nanjing in the northern regions of Yangtze River and Zijinshan National Forest Park of Nanjing in the southern regions of Yangtze River. The continuous large-scale afforestation projects year by year increased the vegetation coverage of these regions, improved the regional microclimate, relieved the heat island effect which was gathered in Nanjing to a small extent, thus made the main urban areas with heat island intensity ascending have the obvious “cold islands”. Aiming at the regions of heat island intensity descending obviously in the northern regions of Yangtze River, this research analyzed and found that obvious descent was due to the ecological restoration and protective development made by the Yizheng Longshan Scenic Spot after being awarded the provincial-level forest park in February 2015, which improved the ecological environment and regional microclimate of these regions, thus further reduced the heat island effect of the surrounding regions.

## Conclusions and discussions

1. Nanjing was a typical quick urbanization place, whose land surface thermal environment pattern showed the below features: in 2014, the northern regions of Nanjing had higher land surface temperature, and the administrative centers of various districts and minor industrial zones formed many scattered land surface temperature high-value regions; in 2016, the west-central regions of Nanjing had higher land surface temperature, and the land surface thermal environment started showing the strip distribution along main roads; in 2018, the land surface thermal environment started showing the distribution features of concentrated block masses, the high-value regions had more concentrated distribution mainly around the main urban areas in the southern regions of Yangtze River and Jiangbei New Area.
2. In view of spatio-temporal change pattern of Nanjing land surface thermal environment, the changes of Nanjing heat island intensity wholly showed the trend of “ascending in urban areas and riverside regions, and descending in suburbs”. The regions of heat island intensity ascending obviously were concentrated around the main urban areas in the southern regions of Yangtze River, Jiangbei New Area and the upstream and downstream southern bank regions of Yangtze River Nanjing Section. Thereinto, in the northern regions of Yangtze River, the regions of heat island intensity ascending obviously showed the distinct strip distribution, and were divided into two-way extension from northeast to southwest direction. The regions of heat island intensity descending obviously were concentrated around the Donggou Town of Liuhe District and riverside belts on the north of Tangshan. Meanwhile, the whole water regions of Yangtze River Nanjing Section, the Shijiu Lake and Gucheng Lake in the southern regions of Nanjing belonged to the regions of heat island intensity descending obviously.
3. In view of the analysis on reasons of changes in Nanjing land surface thermal environment, the changes of administrative division, layout of transportation trunk lines, transfer of industrial centers, ecological construction projects were the important drive factors for evolution of land surface thermal environment pattern of these regions. The cancellation of counties and establishment of districts of Gaochun and Lishui caused two new heat islands appearing in the outer suburbs in the southern Nanjing; the quick development of the regions around the Nanjing Around-city Expressway Jiangbei Section and Jiangbei Avenue Expressway promoted the linear gathering of land surface thermal environment; the construction of Nanjing Jiangbei New Materials High-Tech Park and the projects of closure and relocation of chemical pollution enterprises caused the transfer of urban industrial centers and the obvious changes in the heat environment pattern; the ecological restoration and protective development of Laoshan National Forest Park of Nanjing, Zijinshan National Forest Park of Nanjing, and Yizheng Longshan Scenic Spot prompted the surrounding regions forming cold islands.
4. Following the development of eastern coastal economic belts and deep-going of national significant strategies including Yangtze River Economic Belt Strategy and Yangtze River Delta Integration Strategy, the urbanization levels of Nanjing will keep in the high-level stable stage, and the urban land surface thermal environment pattern will continue gathering, and the severe urban ecology problems will be caused. Therefore, during the process of urban construction, the regional functions should be organized scientifically, and the administrative regions should be divided reasonably; aiming at the potential “heat islands” gathering regions, the ecological construction projects should be arranged in advance in order to timely prevent the forming of “heat islands”; the industrial development orientation should be adjusted and the industrial spatial layout should be optimized in order to relieve the “heat islands”; the ecological project means and ecological protection policies should be used fully so as to lead the forming of “cold islands” and create the excellent urban spaces.

## Supporting information

**S1 Table. Field investigation records.**

(DOCX)

## Acknowledgments

This research was funded by National Natural Science Foundation of China (Grant number 42071282), Foreign Cooperation Projects of Chinese Academy of Sciences (Strategic Layout and Development Guide for World Network of Biosphere Reserves in China).

